# Proteomic and Metabolomic Profiling of Transgenic Pod Borer-Resistant Cowpea: Assessing Unintended Molecular Changes and Their Implications for Ecosystem Resilience

**DOI:** 10.64898/2026.06.24.734197

**Authors:** Abraham Isah, Rebeccah W. Ndana, Malann David Yoila, Abdulrazak Baba Ibrahim, Oluwasijibomi Charles Ogunremi

## Abstract

**Background:** The commercialization of Nigeria’s single-line pod borer-resistant (PBR) cowpea (IT97KT), the first transgenic cowpea variety in the world expressing *Cry1Ab* gene, has raised questions about potential unintended molecular changes and their ecological implications. This study employed integrated proteomic and metabolomic profiling to compare the transgenic line with its non-transgenic isoline (IT97KN) and assess molecular indicators associated with ecosystem resilience.

**Methods:** Proteomic analyses were conducted using LC-MS/MS following filter-assisted sample preparation, while metabolomic profiling employed GC-MS and UHPLC-MS/MS platforms. Differential protein and metabolite abundance were assessed using label-free quantification, volcano plot analysis, principal component analysis (PCA), hierarchical clustering, and Gene Ontology (GO) enrichment analyses.

**Results:** Proteomic profiling revealed substantial overlap between IT97KT and IT97KN, with only a limited subset of proteins exhibiting significant differential abundance. Upregulated proteins in IT97KT were primarily associated with seed storage, redox regulation, oxidative stress mitigation, and defense-related functions, including Late Embryogenesis Abundant Protein 1 (LEA1), vicilins, thioredoxin, and iron superoxide dismutase. Among 37 proteins linked to ecological adaptation, only LEA1, CPRD22, and Bg7S showed significant differences. Similarly, only carbonic anhydrase II displayed differential abundance among proteins associated with potential ecological risk. PCA and clustering analyses demonstrated high proteomic similarity between genotypes. Metabolomic analyses identified sixteen major metabolites, predominantly fatty acids, with no statistically significant differences in abundance or composition between transgenic and non-transgenic lines

**Conclusions:** The transgenic PBR cowpea exhibited minimal unintended proteomic and metabolomic alterations relative to its non-transgenic isoline. These findings indicate that *Cry1Ab* insertion did not substantially disrupt molecular pathways associated with ecological adaptation, environmental risk, or metabolic homeostasis, providing molecular evidence supporting the environmental and biosafety equivalence of PBR cowpea.

## 1.0 Introduction

The global demand for sustainable agriculture has intensified efforts to develop resilient crop varieties that address pest pressures while minimizing environmental impacts (Yuvarajan et al., 2026; Bilal et al.,2026; Wang, 2025; Isah et al., 2025; Isah et al., 2024b; Isah et al., 2024d). In Africa, food security faces challenges from climate change, pest infestations, and declining soil fertility, driving the adoption of genetically modified (GM) crops to enhance productivity and nutritional access (Pawlak & Kołodziejczak, 2020; Tilman et al., 2014; Isah et al., 2024a). Nigeria’s pod borer-resistant (PBR) cowpea (*Vigna unguiculata*), the first transgenic food crop commercialized in the region, incorporates the Cry1Ab gene from *Bacillus thuringiensis* to confer resistance against the devastating *Maruca vitrata* pod borer (Gbashi et al., 2021; Isah et al., 2024a; 2024b). This innovation has increased yields and reduced reliance on chemical pesticides, yet concerns persist about unintended molecular changes and their ecological consequences, particularly for non-target organisms and ecosystem stability (Isah et al., 2024a; 2024c; Fernandes et al., 2022; Lucht, 2015).

Proteometaboiomics, the integrated study of proteins and metabolites, offers a robust framework to uncover unintended molecular changes in transgenic organisms, revealing alterations in biochemical pathways that may influence ecological interactions (Braatz et al., 2021; Nadine et al., 2020). While prior studies have demonstrated that PBR cowpea maintains biodiversity comparable to its non-transgenic isoline (Isah et al., 2024), the molecular basis of these findings remains underexplored. Unintended changes in protein and metabolite profiles could alter plant physiology, stress responses, or interactions with herbivores, pollinators, and soil microbiota, potentially affecting ecosystem resilience (Isah et al., 2024c; Isah et al., 2024d; Romeis et al., 2014; O’Callaghan et al., 2005). For instance, proteomic shifts may influence inducible defense pathways, mirroring stress-responsive mechanisms in long-lived species, while metabolic changes could impact nutrient cycling or trophic interactions (Lopez-Giolar et al., 2023; Schorling & Freier, 2006). In African agroecosystems, where cowpea supports diverse ecological niches and serves as a critical protein source, such changes could have cascading effects on biodiversity and agricultural sustainability (Adom et al., 2019).

This study aims to elucidate the unintended molecular changes in transgenic pod borer-resistant cowpea (IT97KT) compared to its non-transgenic isoline (IT97KN) through comprehensive proteometabolomic profiling, using advanced analytical platforms including GC-MS, UHPLC-MS/MS and LC-MS/MS, to assess their ecological implications and inform sustainable transgenic crop deployment.

### 1.1 Methods

#### Seed Collection and Processing

The Institute for Agricultural Research (IAR) Zaria supplied seeds of the transgenic PBR cowpea (IT97KT) and its isoline, IT97KN. Using the lateral flow strip kits acquired from Qiagen Inc. at Ahmadu Bello University’s Mary Halaway Laboratory, Department of Biochemistry, Faculty of Life Sciences, the Cry1Ab event in the PBR cowpea was verified: The extraction buffer was then added to each container after 5 g of transgenic and non-transgenic seeds were mashed independently in two separate mortars and pestles. The lines were then read after the flow strip was introduced and left there for roughly ten minutes (Isah et al., 2025; Isah et al., 2024a; 2024b; 2024c).

#### Protein Extraction

Ground samples (0.05 g) of transgenic pod borer-resistant (PBR) cowpea (IT97KT) and its non-transgenic isoline (IT97KN) were transported to the Proteomics and Mass Spectrometry Facility (PAMS), University of Georgia for both the proteomics and metabolomics analysis. Proteins were extracted using two buffers: Tris-EDTA (50 mM Tris, 1 mM EDTA, pH 8.0) and LDS (2% SDS, 0.25 M Tris, 1 mM EDTA, 10% glycerol, pH 8.0), selected for their efficacy in plant protein solubilization (Wang et al., 2008). Each sample was mixed with 1 mL of extraction buffer (four vials total), vortexed for 30 s, and subjected to ultrasonication at 37°C for 30 min to enhance protein yield (Isaacson et al., 2006). The process was repeated, followed by centrifugation at 17,000 x g for 30 min. Supernatants were collected and stored at -20°C. For precipitation, 100 uL of supernatant was mixed with 900 uL of cold ethanol (-20°℃), vortexed briefly, and incubated at -20°C for 16 hr to concentrate proteins (Jiang et al., 2004)

#### Filter-Assisted Sample Preparation (FASP)

Proteins were processed using a modified Filter-Assisted Sample Preparation (FASP) method adapted from Nagaraj et al. (2008). Tris-extracted proteins were resuspended in 100 uL of urea buffer (8 M urea, 0.2% sodium deoxycholate, 1M ammonium bicarbonate), while LDS-extracted proteins (50 uL, undiluted due to precipitation failure) were used directly. Proteins were denatured with 5 uL of 0.5 M dithiothreitol (aqueous) at 100°C for 5 min, cooled to room temperature, and alkylated with 10 uL of 0.5 M iodoacetamide (aqueous) to prevent disulfide bond reformation (Shevchenko et al., 2006).

Samples (50 uL) were transferred to 10 kDa MWCO filters (Microcon-10kDa, Millipore). LDS samples were diluted with 400 uL of urea buffer and centrifuged at 14,000 x g for 30 min to remove LDS, which is incompatible with downstream analysis, followed by washes with 200 uL of 0.5x urea buffer and 200 uL of digest buffer (50 mM ammonium bicarbonate, 0.2% sodium deoxycholate) (Wiśniewski et al., 2009). Trypsin (2 ug in 20 mM tetraethylamine) was added, and samples were digested overnight at room temperature (Isah et al., 2024b).

The following day, filters were centrifuged at 14,000 x g for 15 min to collect tryptic peptides (∼200 uL). Peptides were acidified with 2% trifluoroacetic acid, transferred to 1.5 mL Eppendorf tubes, and mixed with an equal volume of ethyl acetate to remove sodium deoxycholate, with the organic layer discarded and the process repeated (Masuda et al., 2008). Peptides were dried using a vacuum concentrator.

#### Liquid Chromatography-Mass Spectrometry (LC-MS/MS) Analysis

Mass spectrometry was performed using a Thermo Fisher LTQ Orbitrap Elite Mass Spectrometer coupled with a Proxeon Easy NanoLC system (Waltham, MA) at PAMS, University of Georgia. Tryptic peptides were resuspended in 100 uL of 0.1% formic acid/2% acetonitrile, and 1 uL was loaded onto a 13-cm x 100 um i.d. column packed with Dr. Maisch ReproSil-Pur 120 C18 AQ resin (3 um; Ammerbuch, Germany). A 105-min gradient elution was run using buffers A (0.1% formic acid) and B (0.1% formic acid in acetonitrile), increasing from 2% to 50% B over 95 min at 300 nL/min, then to 95% B over 10 min, optimized for peptide separation (Thakur et al., 2011). Peptides were directly eluted into the mass spectrometer. Data were acquired using Xcalibur software (Version 3.0; Thermo Fisher Scientific) in data-dependent acquisition (DDA) mode, with a survey MS scan followed by CID MS/MS analysis of the top 10 ions. MS and MS/MS scans were acquired at resolutions of 120,000 and 15,000, respectively, ensuring high mass accuracy (Olsen et al., 2007).

#### Protein Identification and Quantification

Protein identification and modification characterization were performed using Thermo Fisher Proteome Discoverer (Version 3.0 SP1) with SEQUEST HT, referencing the Vigna unguiculata database (NCBI, txid3917, 82,536 entries, March 2023). Search parameters included a precursor ion m/z tolerance of 10 ppm, product ion m/z tolerance of 0.02 Da, maximum of two missed cleavages, dynamic modification (oxidation on methionine), and static modification (carbamidomethyl on cysteine). Results were validated using Percolator with a concatenated decoy database, targeting false discovery rates of 0.01 (strict) and 0.05 (relaxed) (Kall et al., 2007). Semi-quantitative analysis was conducted using a label-free quantification workflow in Proteome Discoverer, summing the intensities of unique and shared (razor) peptides to estimate protein abundance (Cox and Mann, 2008). Ratios of IT97KN/IT97KT from Tris and LDS extractions were compared to identify differentially expressed proteins, visualized in volcano plots with - log10 p-value (y-axis) and log2 ratio (x-axis). Proteins were classified as up- or down-regulated based on at least two unique peptides, a log2 ratio >2 or <- 2, and consistent changes across Tris and LDS datasets, following established criteria for differential expression (Tyanova et al., 2016). Results were exported to spreadsheets.

#### Metabolomic Profiling

Metabolomic analysis of transgenic pod borer-resistant (PBR) cowpea (IT97KT) and its non-transgenic isoline (IT97KN) was performed at the Proteomics and Mass Spectrometry Facility (PAMS), University of Georgia, using three complementary platforms: ultrahigh-performance liquid chromatography/tandem mass spectrometry (UHPLC-MS/MS) optimized for basic species, UHPLC-MS/MS optimized for acidic species, and gas chromatography/mass spectrometry (GC-MS). Ground cowpea samples (0.05 g) were extracted in 1 mL of 80% methanol with 0.1% formic acid, vortexed for 30 s, and sonicated at 37°C for 15 min to maximize metabolite recovery (De Vos et al., 2007). Extracts were centrifuged at 17,000 x g for 10 min, and supernatants were divided into three equal aliquots for analysis on each platform, following established protocols for comprehensive metabolomic profiling (Fiehn et al., 2008).

For UHPLC-MS/MS, analyses were conducted using a Thermo Fisher Q Exactive Plus mass spectrometer coupled with a Vanquish UHPLC system (Waltham, MA). Chromatographic separation was performed on a Waters ACQUITY UPLC BEH C18 column (2.1 mm × 100 mm, 1.7 um) at a flow rate of 0.35 mL/min. For basic species, mobile phases consisted of A (0.1% formic acid in water) and B (0.1% formic acid in acetonitrile), with a gradient from 5% to 95% B over 12 min. For acidic species, mobile phase A was 5 mM ammonium acetate (pH 8.0), and B was acetonitrile, with a similar gradient (Patti et al., 2012). Full-scan mass spectra (m/z 100-1000) were acquired at 70,000 resolution, followed by data-dependent MS/MS of the top 10 ions, with collision energies of 10, 20, and 30 eV (Want et al., 2010).

For GC-MS, samples were derivatized with 50 uL of N.O-bis(trimethylsilyl)trifluoroacetamide (BSTFA) containing 1% trimethylchlorosilane at 70℃ for 30 min (Fiehn, 2006). Analysis was performed on an Agilent 7890A GC system coupled with a 5975C MSD (Santa Clara, CA), using a DB-5MS column (30 m × 0.25 mm, 0.25 um). The temperature gradient started at 60C, increased to 325C at 10C/min, and held for 5 min, with helium as the carrier gas at 1 mL/min. Mass spectra were acquired in electron ionization mode (70 eV, m/z 50-600).

Metabolites were identified by comparing retention times, molecular weights (m/z), preferred adducts, in-source fragments, and MS/MS spectra to an in-house reference library of chemical standards, supplemented by the NIST 17 Mass Spectral Library (Kind et al., 2009) Data were processed using Thermo Xcalibur (Version 3.0) for UHPLC-MS/MS and Agilent MassHunter for GC-MS. Label-free quantification was performed by integrating peak areas of precursor ions, normalized to total ion current, to compare metabolite abundances between IT97KT and IT97KN (Dunn et al., 2011). Differential metabolites were identified using volcano plots, with significance thresholds of log2 fold change >1 or <- 1 and -log10 p-value >1.3, analyzed in R (Version 4.2.1) with the MetaboAnalyst package (Chong et al., 2018).

## 2.0 Results

### 2.1 Proteomic Profiling Reveals Limited Differential Protein Abundance Between Transgenic and Non-Transgenic Cowpea

LC-MS/MS-based proteomic analysis was conducted to evaluate potential unintended molecular changes between the transgenic pod borer-resistant cowpea (IT97KT) and its non-transgenic isoline (IT97KN). Proteins extracted using both Tris and LDS buffers were subjected to label-free quantitative analysis.

Volcano plot analysis revealed a relatively small subset of proteins exhibiting significant differential abundance between the two cowpea lines (**Figure 1**). Although both extraction methods identified genotype-specific proteins, the Tris extraction protocol recovered a greater number of significantly altered proteins than the LDS protocol. Differentially abundant proteins were detected in both genotypes, indicating localized molecular variation rather than widespread proteomic disruption.

**Figure 1:**
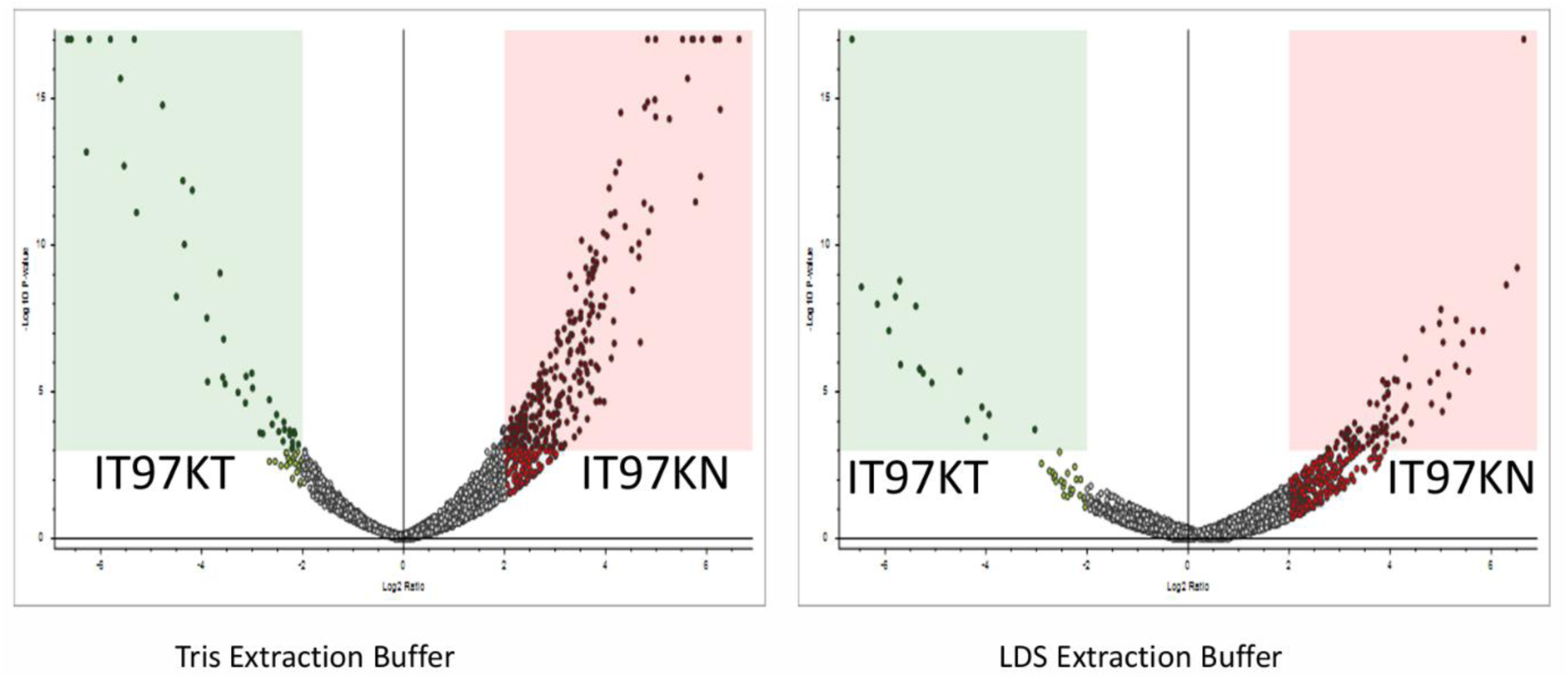
LC/MS/MS Volcano Plots Comparison of the Biological Peptides

Overall, the distribution of proteins across both datasets suggested substantial overlap between the proteomes of IT97KT and IT97KN, with only a limited number of proteins showing statistically significant abundance changes.

### 2.2 Differentially Abundant Proteins Are Primarily Associated With Storage, Stress Response and Redox Functions

A total of several proteins were identified as significantly upregulated in either IT97KT or IT97KN (**Tables 1a and 1b**). Among the proteins enriched in IT97KT were Late Embryogenesis Abundant Protein 1 (LEA1), vicilin and vicilin-like storage proteins, β-conglycinin subunit 1-like protein, thioredoxin 1, formate dehydrogenase 1, iron superoxide dismutase, and actin-11.

**Table 1.**
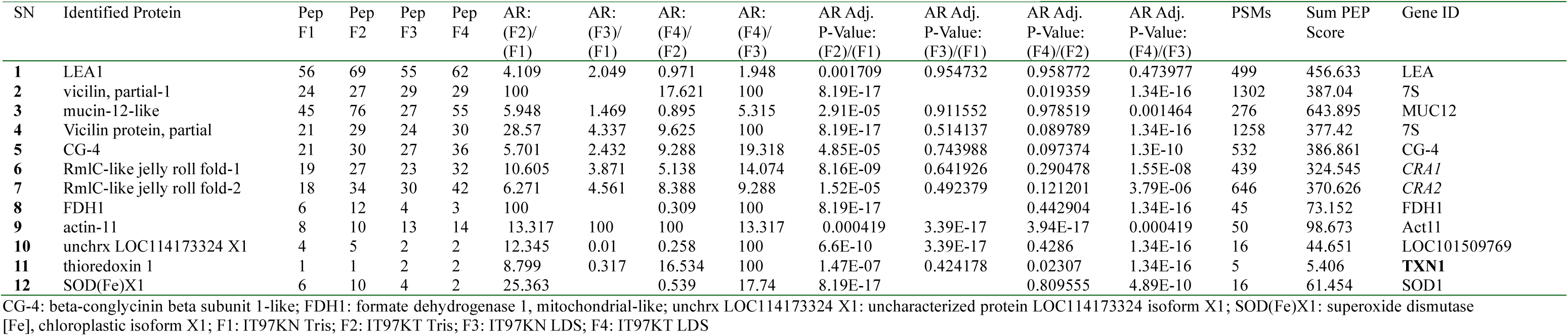
a: Up regulated proteins in both transgenic IT97KT and its conventional non transgenic IT97KN cowpea samples.

**Table 1.**
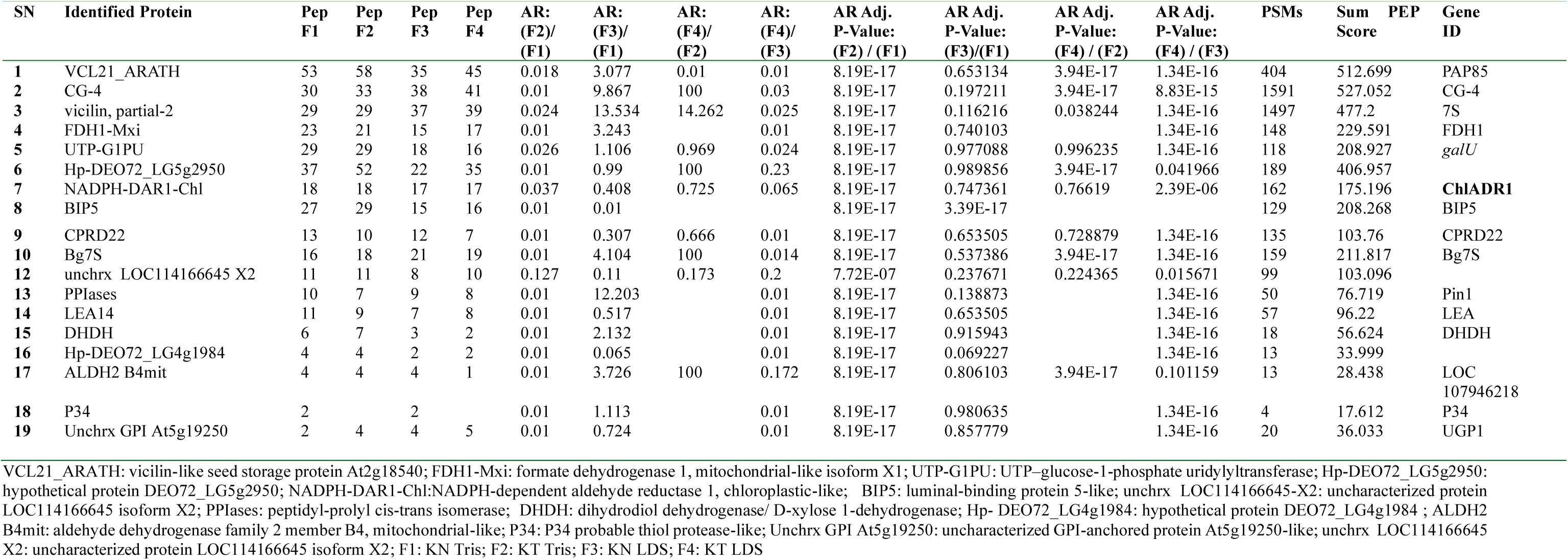
b: Down regulated proteins in the two cowpea samples.

Many of the proteins showing increased abundance in IT97KT are involved in seed storage, oxidative stress mitigation, cellular redox regulation, and energy metabolism. Conversely, proteins exhibiting reduced abundance in IT97KT were predominantly associated with carbohydrate metabolism, stress adaptation, and energy reserve mobilization.

Despite these differences, the majority of identified proteins did not display statistically significant changes, indicating that transformation did not substantially alter the overall proteomic composition of the cowpea seed.

### 2.3 Ecological Adaptation-Associated Proteins Exhibit Minimal Alteration

To explore molecular indicators associated with ecological fitness and environmental adaptation, proteins previously associated with stress tolerance and adaptive responses were examined.

Thirty-seven proteins linked to ecological adaptation processes were identified (Table 2). Of these, only three proteins met the criteria for significant differential abundance. LEA1 was significantly upregulated in IT97KT, whereas CPRD22 and Bg7S were significantly downregulated.

**Table 2:**
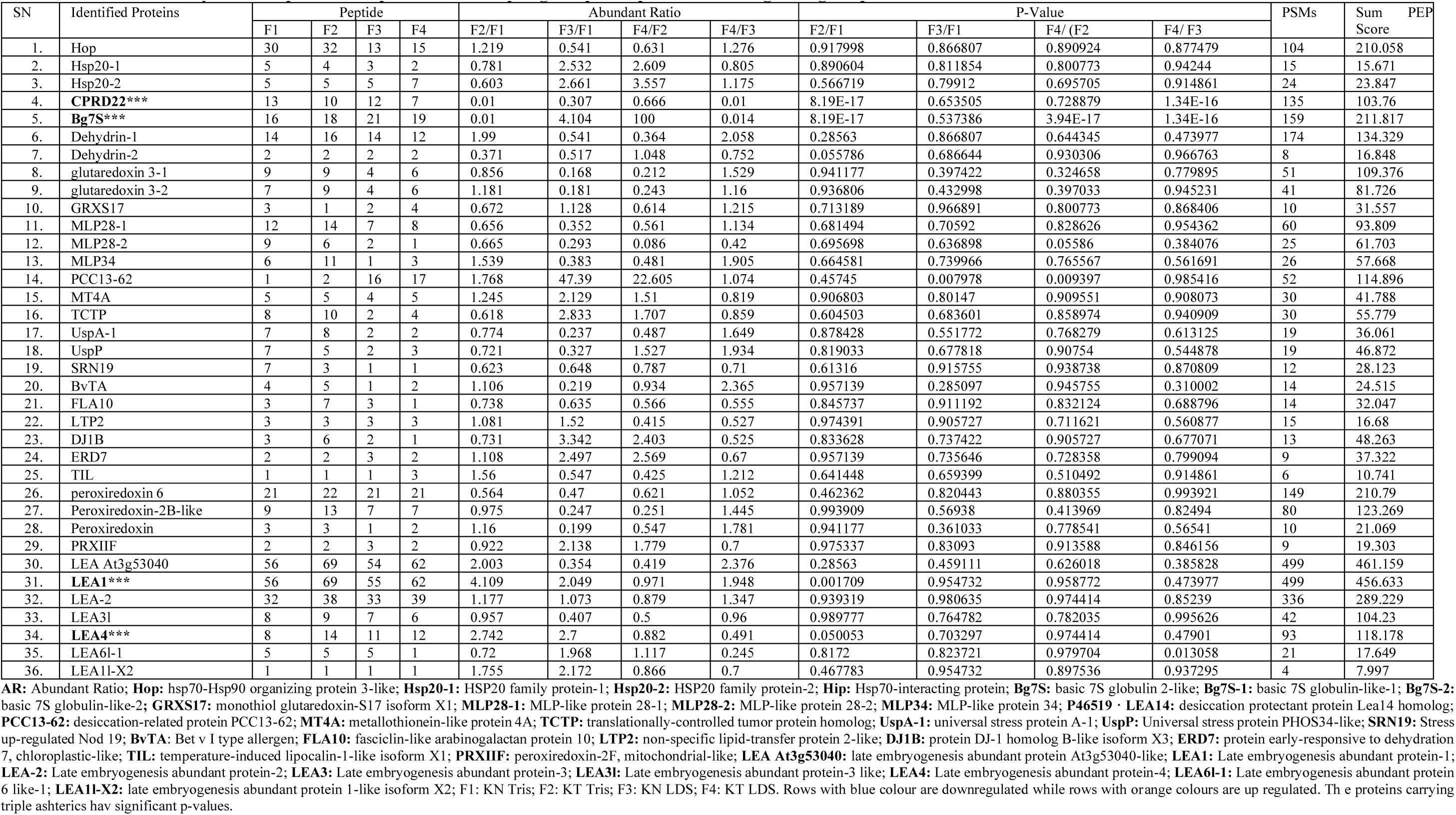
Differential analysis of the proteins responsible for adapting cowpea to possible evolving ecological patterns.

Most adaptation-related proteins, including dehydrins, heat shock proteins, lipid transfer proteins, glutaredoxins, universal stress proteins, and peroxiredoxins, showed no statistically significant differences between the transgenic and non-transgenic lines. These findings indicate that the genetic modification had minimal influence on proteins associated with environmental adaptation and stress resilience.

### 2.4 Potential Ecological Risk-Associated Proteins Remain Largely Unchanged

Nine proteins previously implicated in ecological interactions and environmental responses were evaluated (**Table 3**).

**Table 3:** Proteins Associated with Ecological Risk Assessment.

|  |  | Pep |  |  |  | Abundant Ratio |  |  |  | P-Value |  |  |  | PS<br>Ms | Sum<br>PEP<br>Score | Gene<br>ID |
| --- | --- | --- | --- | --- | --- | --- | --- | --- | --- | --- | --- | --- | --- | --- | --- | --- |
|  | Protein | F1 | F2 | F3 | F4 | F2/F1 | F3/F1 | F4/F2 | F4/F3 | F2/F1 | F3/F1 | F4/F2 | F4/F3 |  |  |  |
| 1 | CA II | 4 | 1 | 5 | 2 | 0.323 | 1.523 | 2.785 | 0.473 | 0.02181 | 0.9060 | 0.6730 | 0.4355 | 12 | 42.828 | **CA2** |
| 2 | PR2 | 13 | 16 | 9 | 9 | 1.498 | 0.64 | 0.791 | 1.972 | 0.693481 | 0.911433 | 0.945755 | 0.52918 | 81 | 101.935 | **PR2** |
| 3 | PR1 | 1 | 2 | 1 |  | 0.869 | 1.172 | 1.649 | 0.98 | 0.941177 | 0.996187 | 0.945755 | 0.975924 | 4 | 8.188 | GLIPR2 |
| 4 | defensin,partial | 6 | 7 | 7 | 7 | 2.053 | 0.294 | 0.579 | 2.243 | 0.256239 | 0.637684 | 0.84013 | 0.383918 | 73 | 64.248 | **PDF1.1** |
| 5 | Defensin-like1 | 2 | 2 | 3 | 3 | 1.296 | 5.458 | 3.845 | 0.913 | 0.872765 | 0.4123 | 0.441878 | 0.973183 | 15 | 24.408 | **DEFB1** |
| 6 | Defensin-like2 | 2 | 2 | 4 | 6 | 1.026 | 18.018 | 16.866 | 1.306 | 0.991971 | 0.068924 | 0.021862 | 0.868251 | 28 | 24.318 | **defl2** |
| 7 | osmotin-like<br>protein | 2 | 2 | 3 | 1 | 0.73 | 3.188 | 1.627 | 0.372 | 0.832993 | 0.640323 | 0.958377 | 0.183701 | 8 | 14.681 | PdOLP1 |
| 8 | Gamma<br>Purothionin | 2 | 2 | 2 | 2 | 1.764 | 1.716 | 1.053 | 1.082 | 0.461048 | 0.915943 | 0.996011 | 0.969357 | 9 | 8.69 | THI1.3 |
| 9 | HS1/DABB1-like | 4 | 5 | 3 | 4 | 0.75 | 0.431 | 0.646 | 1.074 | 0.856318 | 0.711511 | 0.84013 | 0.921336 | 9 | 13.273 | **HS1** |

**CA II:** carbonic anhydrase 2-like; **PR2:** pathogenesis-related protein 2; **PR1:** pathogenesis-related protein 1; **Defensin-like1:** *RecName: Full=Defensin-like protein I; AltName: Full=Cp-thionin I; AltName: Full=Cp-thionin-1; AltName: Full=Gamma-thionin I*; **Defensin-like2:** *RecName: Full=Defensin-like protein 2; AltName: Full=Cp-thionin II; AltName: Full=Cp-thionin-2; AltName: Full=Gamma-thionin II*; **HS1/DABB1-like:** *stress-response A/B barrel domain-containing protein HS1-like*

Only carbonic anhydrase II (CA II) exhibited significant differential abundance between the two cowpea lines. All remaining proteins, including defensins, purothionins, osmotin-like proteins, and disease resistance-associated proteins, showed no significant differences.

The limited number of altered proteins within this category suggests that transgene insertion did not substantially alter molecular markers associated with potential ecological risk.

### 2.5 Global Proteomic Profiles Demonstrate High Similarity Between Cowpea Lines

Hierarchical clustering and principal component analyses (PCA) were performed to evaluate overall proteomic variation between IT97KT and IT97KN.

Heatmap clustering revealed similar protein expression patterns across experimental groups, with no distinct separation between transgenic and non-transgenic samples (**Figures 2, 4 and 6**). Likewise, PCA demonstrated strong overlap between sample groups, with the first principal component accounting for more than 90% of the observed variance in most analyses (**Figures 3, 5 and 7**). Collectively, these analyses indicate that the global proteomic architecture of IT97KT remains highly comparable to that of its non-transgenic isoline.

**Figure 2:**
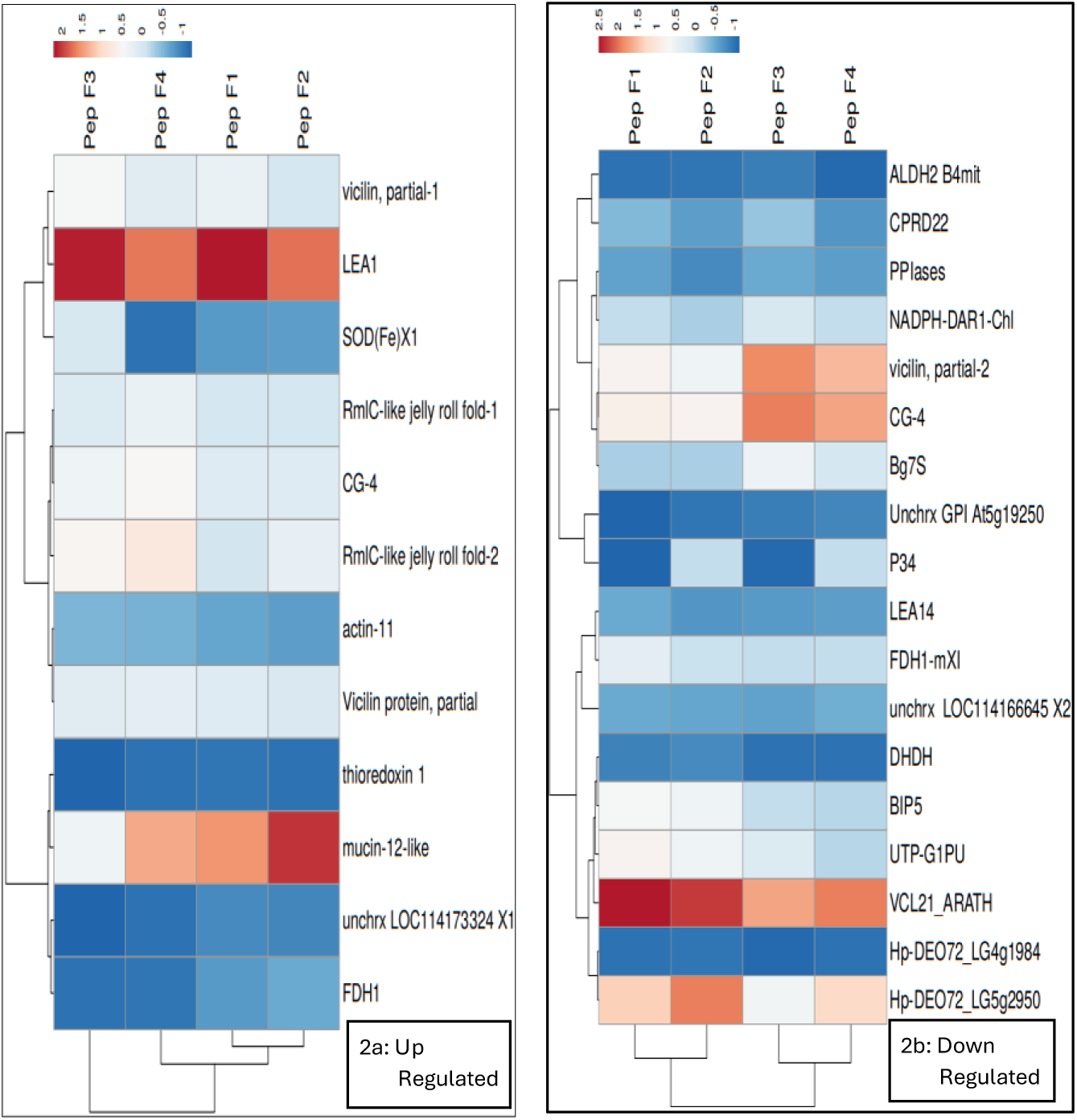
Heat Map of the up regulated protein expression pattern in the transgenic IT97KT cowpea and its isoline, IT97KN under two different buffer conditions (Tris and LDS buffer).

**Figure 3:**
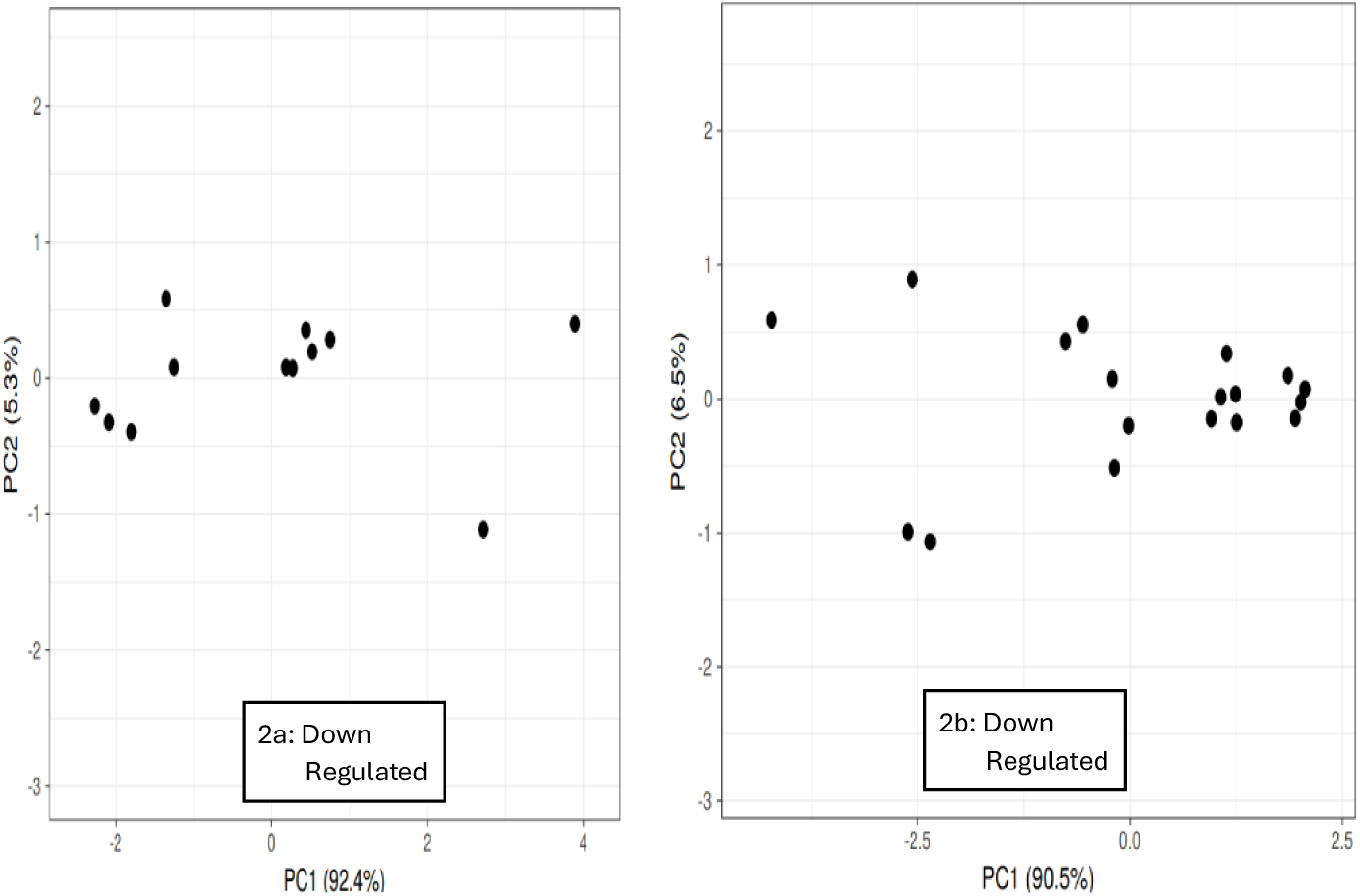
PCA Analysis of the Up and Down Regulated Protein

**Figure 4:**
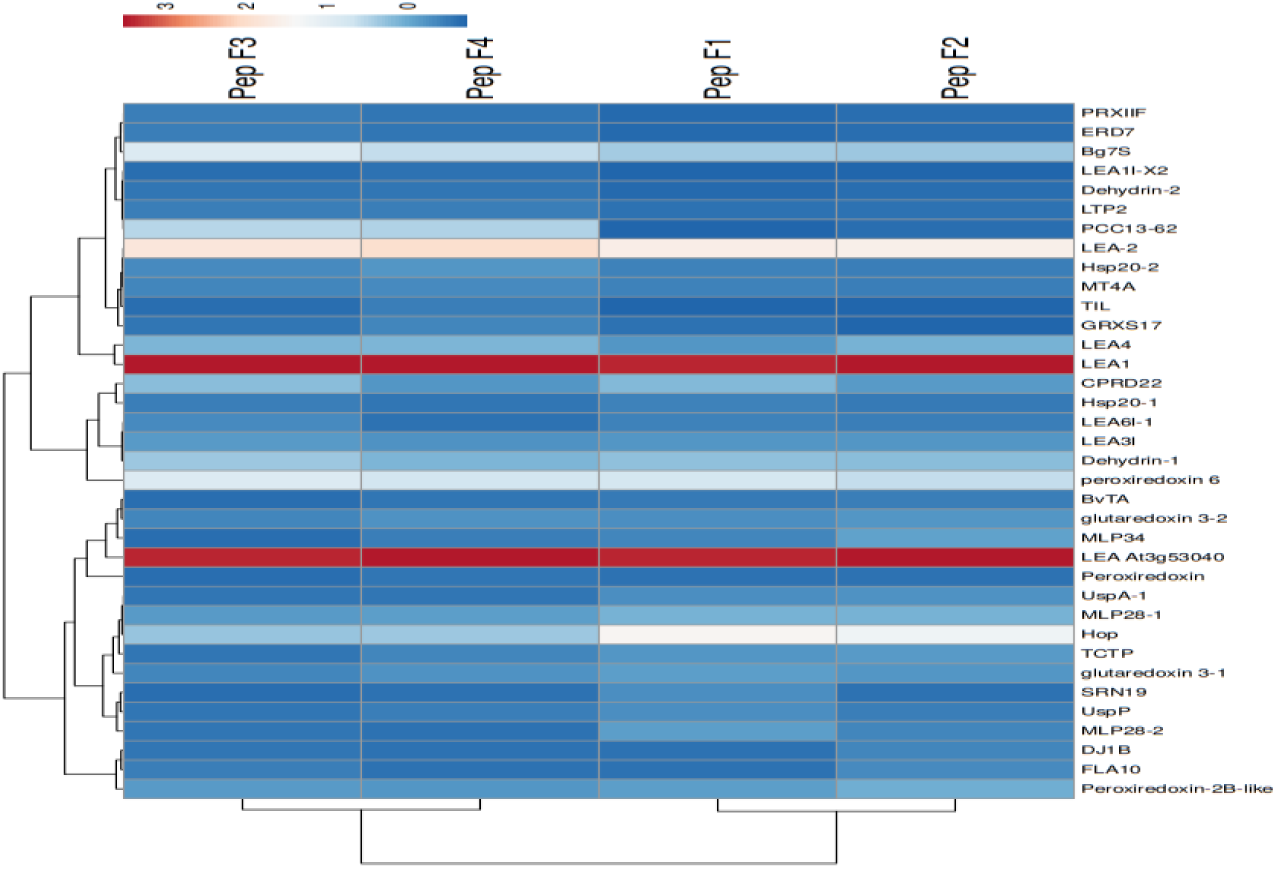
Heat Map of the stress-responsive and adaptation-related proteins

**Figure 5:**
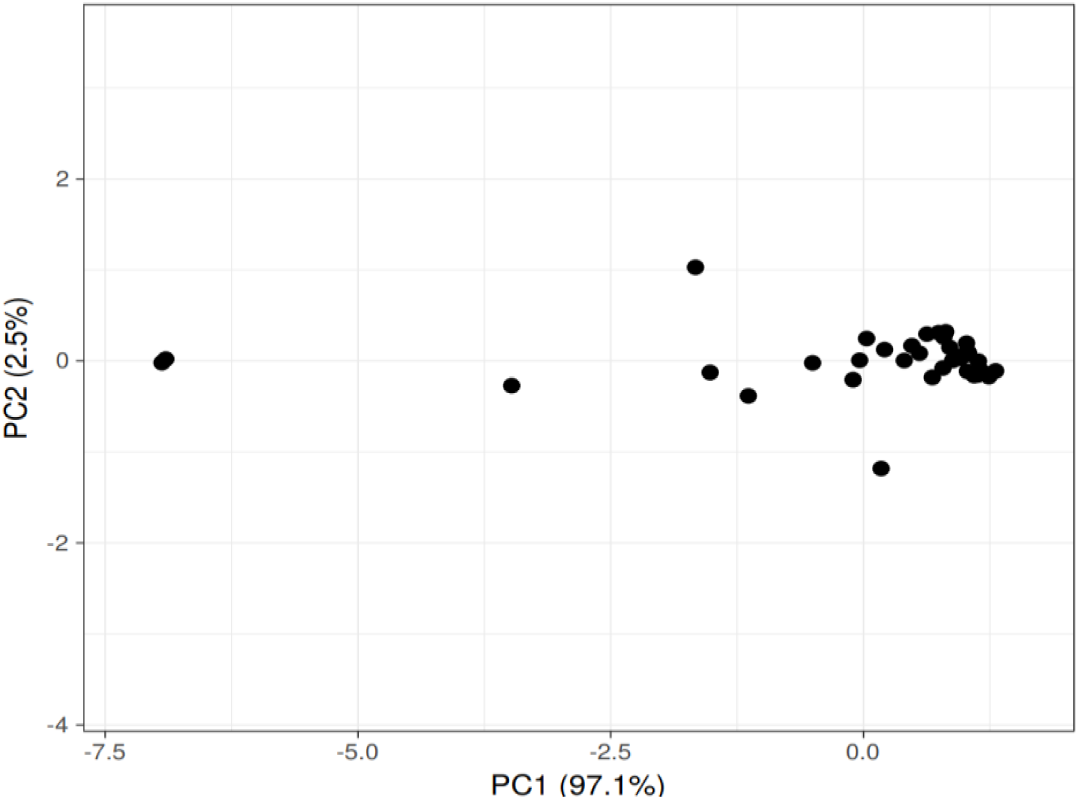
PCA of the Stress-responsive and adaptation-related proteins

**Figure 6:**
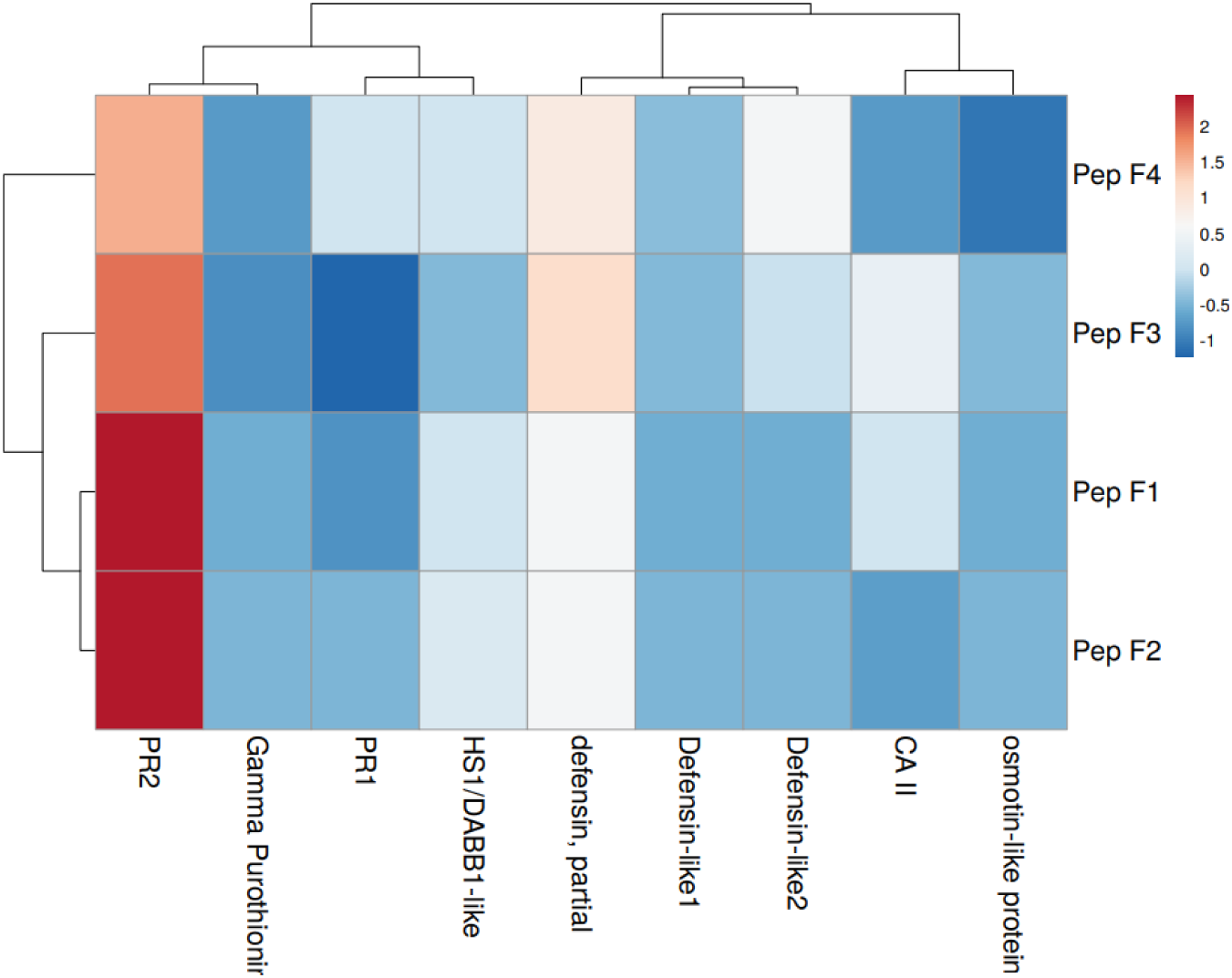
Heatmap of Ecological Risk-Associated Proteins

**Figure 7:**
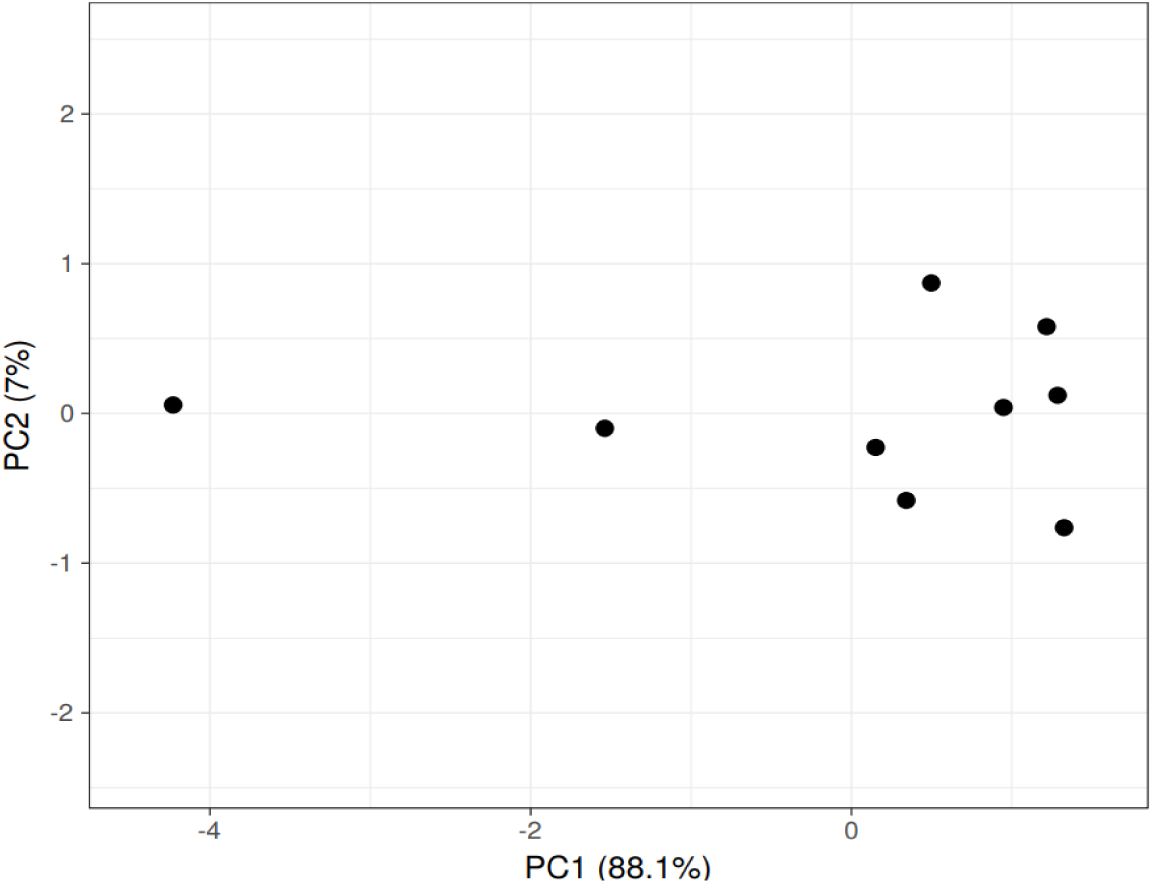
PCA of Ecological Risk-Associated Proteins

#### Clustering Analysis of the Ecological Risk-Associated Proteins

The expression pattern was similar for all the proteins throughout the two different experimental conditions. Exact gradient value was observed for PR2 at between PepF1 and PepF2, Defensin-like 1 at between PepF3 and PepF4, Defensin partial at between Pep F1 and Pep F2 and Gamma Purothionir at between Pep F3 and Pep F4 (**Figure 6**). All other proteins had a close-range value.

The principal component analysis using a unit variance scaling, and the singular value decomposition (svd) shows 88.1% and 7% of total variance in x and y axis (**Figure 7),** consequently showing that the variation among the environmental risk-associated proteins between the transgenic line IT97KT and its isoline, IT97KN is negligible.

### 2.6 Functional Classification and Biological Properties of Differentially Abundant Proteins

#### 2.6.1 Upregulated proteins

##### Functional Classification

Gene Ontology (GO) enrichment analysis revealed that proteins upregulated in IT97KT were primarily associated with biological processes related to pollen tube adhesion, cell-cell adhesion, carbon utilization, defense response to fungi, response to external biotic stimuli, defense response to other organisms, pollen-pistil interaction, and pollen tube development (**Figure 9a**). The highest enrichment factors were observed for pollen tube adhesion, cell-cell adhesion, biological processes involved in intraspecies interaction between organisms, carbon utilization, and defense response to fungus.

**Figure 9.**
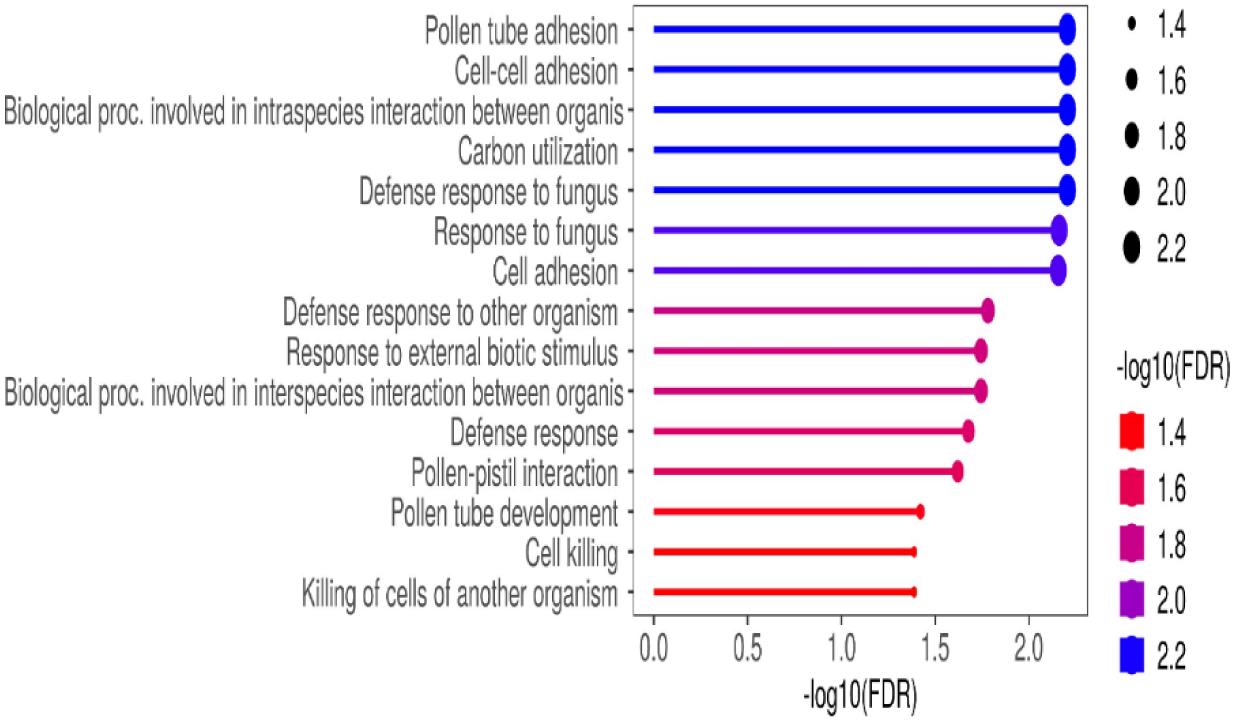
a: GO enrichment of upregulated proteins

**Figure 9.**
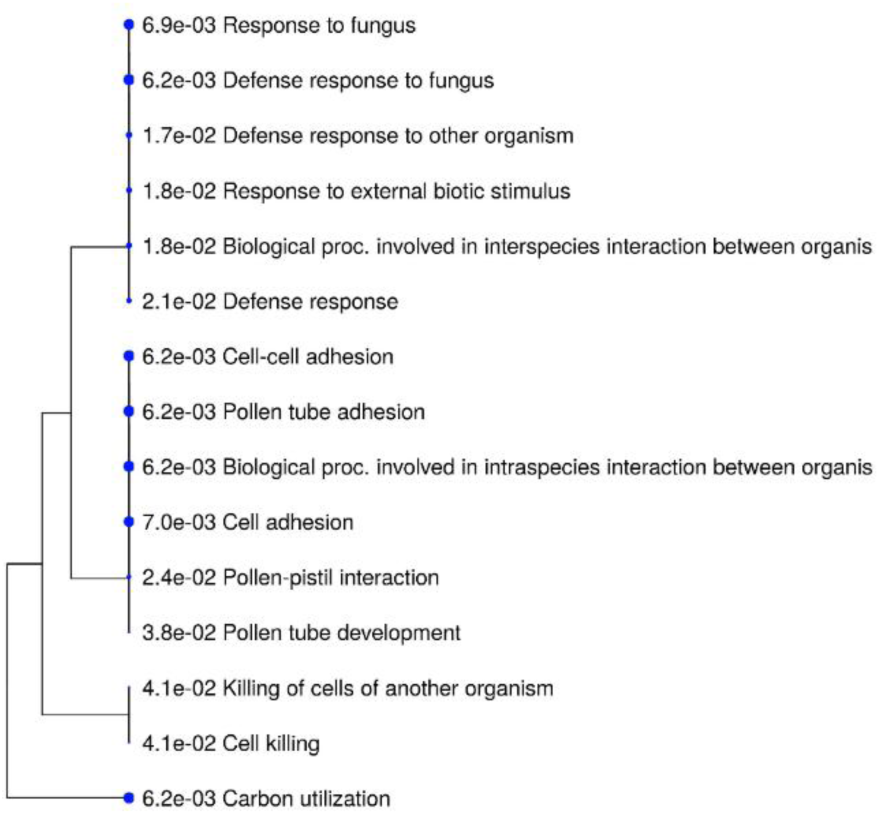
b: A hierarchical clustering tree summarizing the correlation among significant pathways listed in the Enrichment analysis of the Upregulated Proteins

**Figure 9.**
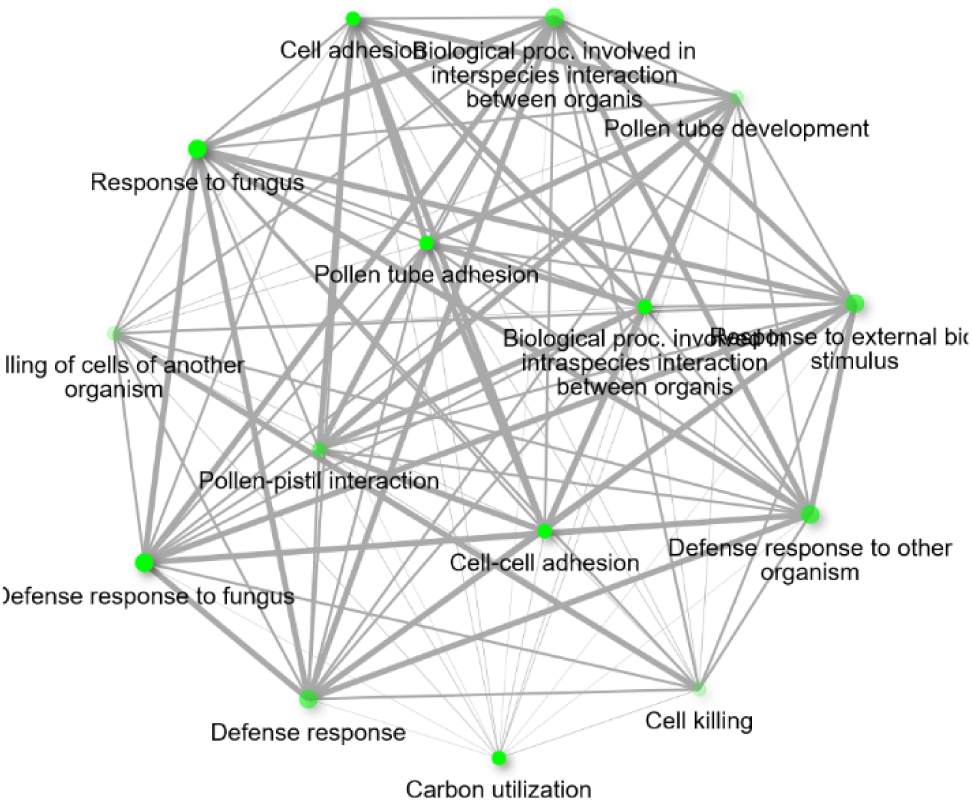
c: Interaction plot of the relationship between enriched pathways of the upregulated proteins

##### Pathway Correlation Analysis

Hierarchical clustering analysis demonstrated strong correlations among several enriched biological processes (**Figure 9b**). Pathways associated with fungal defense responses, defense against other organisms, response to external biotic stimuli, and interspecies interactions formed a closely related cluster. Similarly, cell adhesion-related pathways, including cell-cell adhesion, pollen tube adhesion, pollen-pistil interaction, and pollen tube development, clustered together, indicating shared biological functions among the upregulated proteins.

##### Interaction Network of Enriched Biological Processes

Network analysis revealed substantial overlap among enriched biological processes associated with upregulated proteins (**Figure 9c**). Defense response to fungus and response to fungus displayed the largest and most highly enriched gene sets, as indicated by their larger and darker nodes. Cell-cell adhesion, carbon utilization, and pollen tube adhesion also exhibited strong enrichment despite involving comparatively smaller gene sets. Collectively, these findings suggest coordinated regulation of defense- and adhesion-related biological pathways in the transgenic cowpea line.

#### 2.6.2 Downregulated Proteins

GO enrichment analysis indicated that the downregulated proteins were primarily associated with UDP-glucose metabolic process, glycogen metabolic process, energy reserve metabolic process, callose deposition in cell wall, cell wall thickening, nucleotide-sugar metabolic process, and cold acclimation (**Figure 10a**). Among these biological processes, UDP-glucose metabolic process exhibited the highest enrichment score.

**Figure 10.**
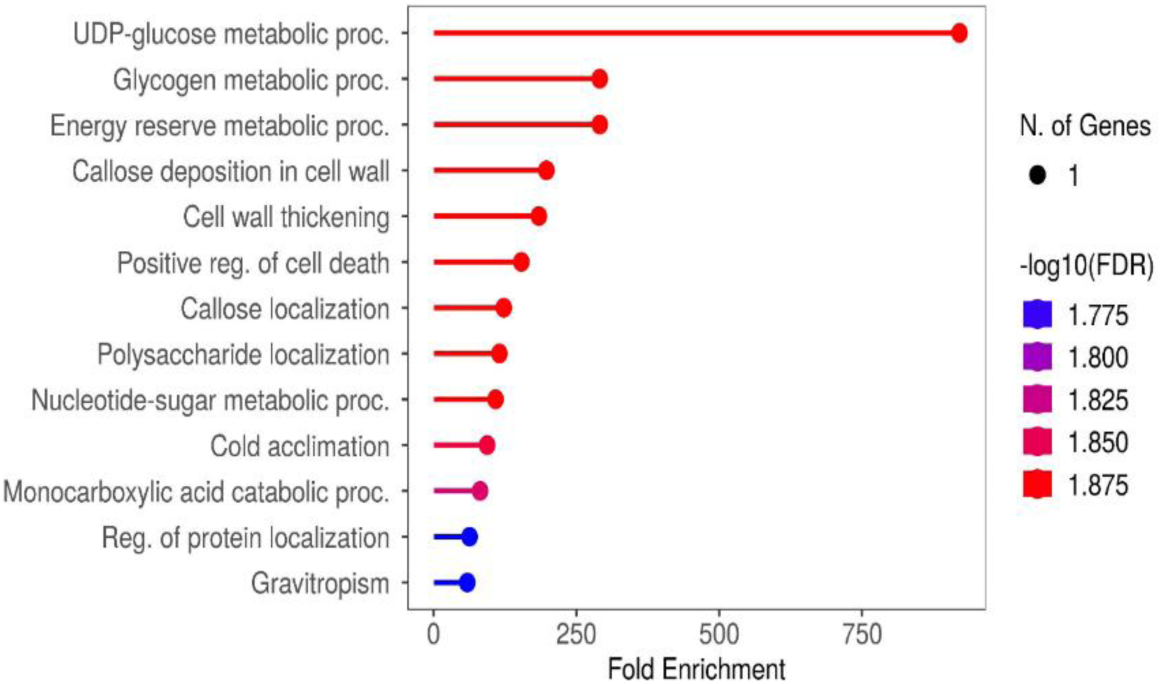
a. GO biological process enrichment analysis of downregulated proteins.

**Figure 10.**
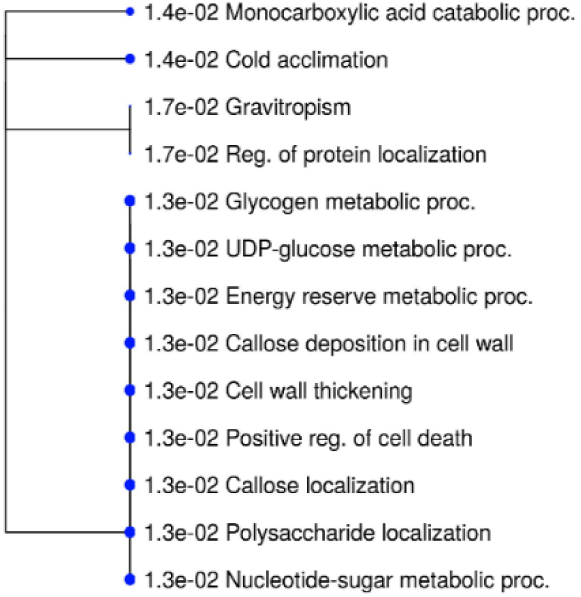
b. Hierarchical clustering of enriched GO biological processes associated with downregulated proteins.

**Figure 10.**
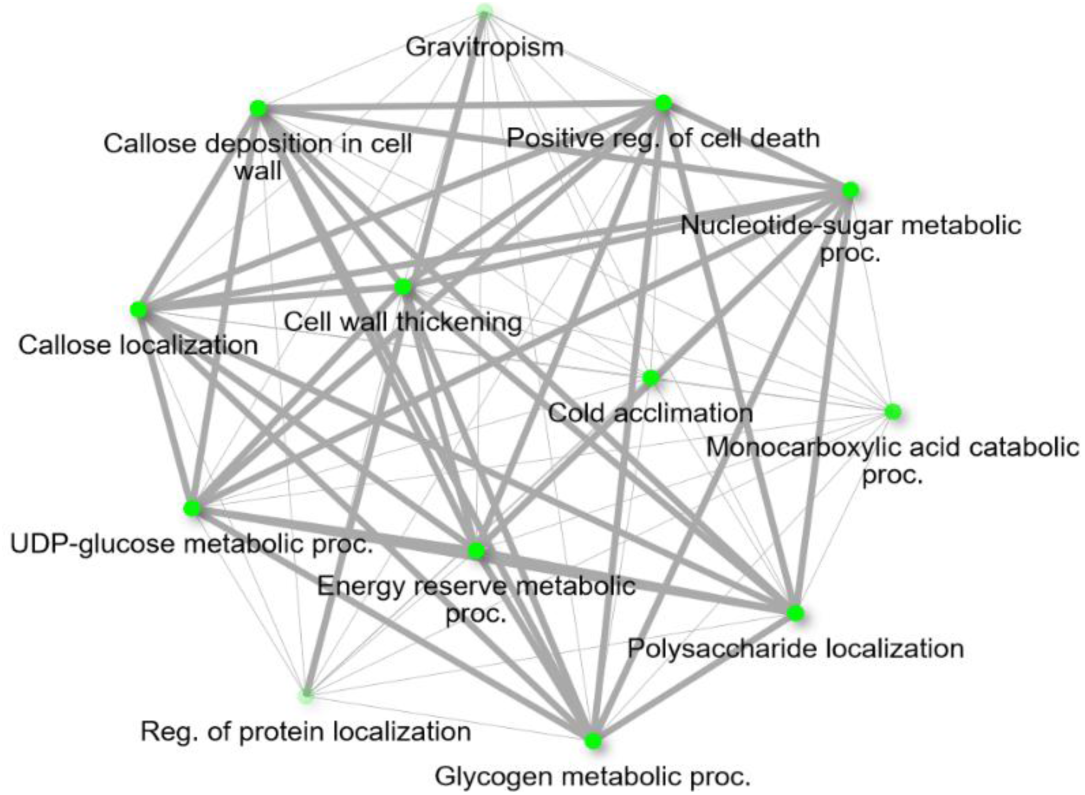
c. Interaction network of enriched GO biological processes associated with downregulated proteins; Two pathways (nodes) are connected if they share 20% (default) or more genes. Darker nodes are more significantly enriched gene sets. Bigger nodes represent larger gene sets. Thicker edges represent more overlapped genes.

##### Pathway Correlation Analysis

Hierarchical clustering analysis revealed strong associations among carbohydrate metabolism- and cell wall-related pathways. Glycogen metabolic process, UDP-glucose metabolic process, energy reserve metabolic process, callose deposition in cell wall, cell wall thickening, polysaccharide localization, and nucleotide-sugar metabolic process clustered closely together, suggesting coordinated biological functions among the downregulated proteins (**Figure 10b**).

##### Pathway Interaction Network

Network analysis further demonstrated extensive overlap among the enriched biological processes. UDP-glucose metabolic process, glycogen metabolic process, callose localization, polysaccharide localization, cold acclimation, nucleotide-sugar metabolic process, positive regulation of cell death, and cell wall thickening were represented by larger and darker nodes, indicating both stronger enrichment and larger associated gene sets. In contrast, regulation of protein localization and gravitropism exhibited smaller and less enriched gene sets (**Figure 10c**).

### 2.7 Metabolomic Profiling Reveals No Evidence of Major Metabolic Perturbation

GC-MS and UHPLC-MS/MS analyses were conducted to determine whether transgene insertion resulted in unintended metabolic alterations.

Chromatographic profiles of IT97KT and IT97KN exhibited highly similar peak distributions and intensities (**Figures 11a & 11b**). A total of sixteen metabolites were identified across both cowpea lines, predominantly consisting of fatty acids including arachidic acid, palmitic acid, α-linolenic acid, oleic acid, stearic acid, and γ-linolenic acid (**Table 4**).

**Figure 11.**
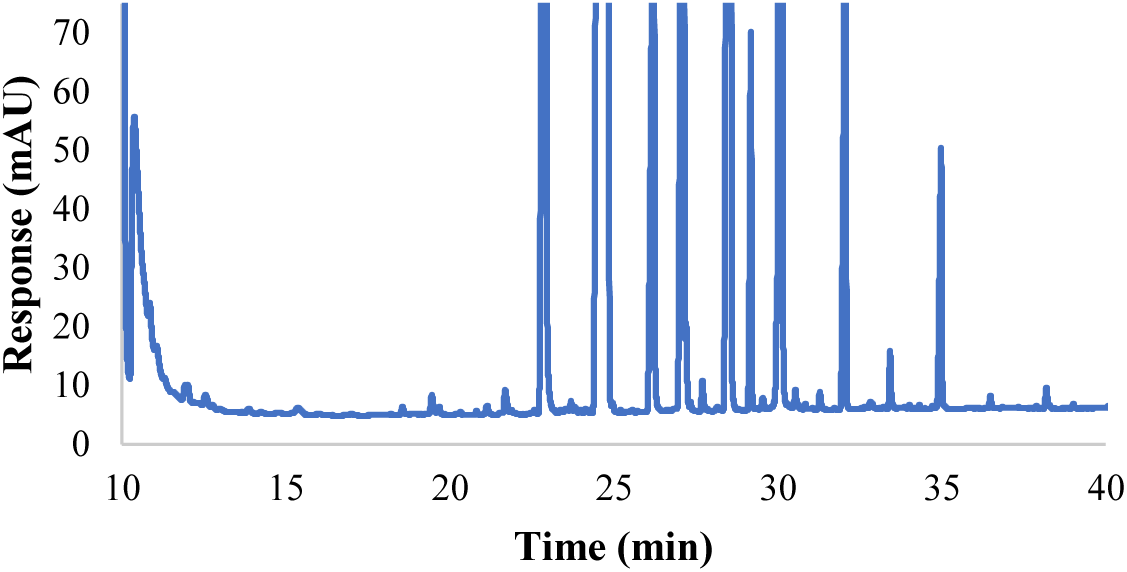
a: Chromatogram of the IT97KN Non-Transgenic Cowpea’s quantified molecular metabolites using the Gas Chromatography-Mass Spectrometry (GC-MS)-based metabolomics method; Zoomed in with a y-axis of 75 units

**Figure 11.**
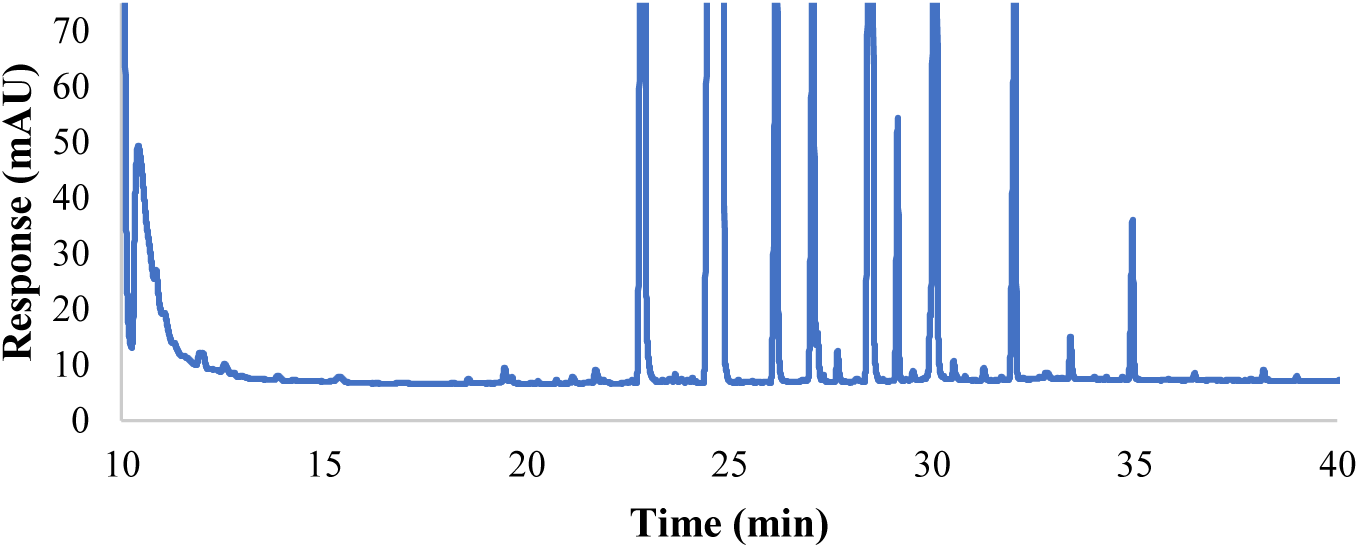
b: Chromatogram of the IT97KT Transgenic Cowpea’s quantified molecular metabolites using the Gas Chromatography-Mass Spectrometry (GC-MS)-based metabolomics method; Zoomed in with a y-axis of 75 units.

**Figure 12:**
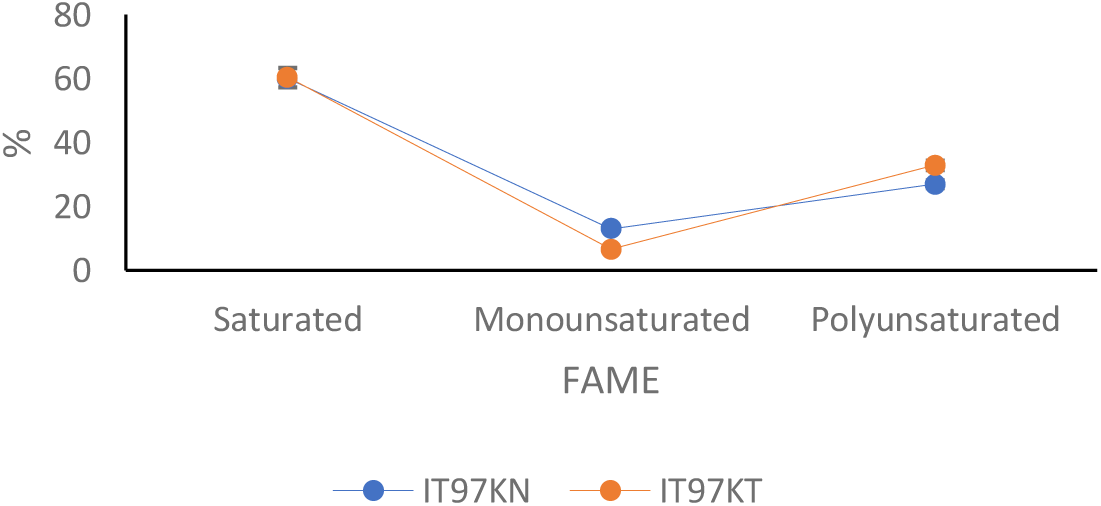
Percentage Analysis of the Saturated, Monosaturated and Polyunsaturated FAME

**Table 4:** Mean Data of the GCMS Chromatogram Analysis.

| Compound | Full names (acid) | Mean |  | P Value | R Square |
| --- | --- | --- | --- | --- | --- |
|  |  | KN | KT |  |  |
| Saturated |  |  |  |  |  |
| C6:0 | Caproic | 0.192±0.002 | 0.252±0.002 | 0.7393 | 0.999 |
| C8:0 | Octanoic | 0.197±0.003 | 0.195±0.003 | 0.5286 | 0.225 |
| C14:0 | Myristic | 0.175±0.002 | 0.218±0.002 | 0.6878 | 0.995 |
| C15:0 | Pentadecanoic | 0.078±0.003 | 0.088±0.003 | 0.6020 | 0.889 |
| C16:0 | Palmitic | 22.568±0.424 | 23.120±0.424 | 0.6415 | 0.552 |
| C18:0 | Stearic | 6.674±0.095 | 6.472±0.095 | 0.5832 | 0.747 |
| C20:0 | Arachidic | 28.055±0.254 | 28.333±0.254 | 0.7251 | 0.425 |
| Monounsaturated |  |  |  |  |  |
| C18:1n-9 | Oleic | 12.874±0.323 | 6.438±0.323 | 0.3116 | 0.995 |
| C18:1n-7 | cis-Vaccenic | 0.435±0.003 | 0.429±0.003 | 0.7393 | 0.714 |
| C20:1n-9 | 9-Eicosenoic | 0.080±0.003 | 0.203±0.003 | 0.4558 | 0.999 |
| Polyunsaturated |  |  |  |  |  |
| C18:3n-6 (GLA) | γ-Linolenic | 0.084±0.058 | 2.247±0.057 | 0.7353 | 0.999 |
| C18:3n-3 (ALA) | α-linolenic | 21.228±0.076 | 23.396±0.076 | 0.7036 | 0.998 |
| C20:2n-6 | Eicosadienoic | 0.123±0.003 | 0.127±0.003 | 0.6858 | 0.587 |
| C18:3n-6 (DGLA) | gamma linolenic | 4.761±0.083 | 6.053±0.085 | 0.4805 | 0.992 |
| C20:4n-6 (AA) | Arachidonic | 0.324±0.095 | 0.364±0.095 | 0.6478 | 0.645 |
| C20:5n-3 (EPA) | eico-sapentaenoic | 2.089±0.323 | 1.722±0.323 | 0.5483 | 0.997 |

where C6:0 - caproic acid; C8:0 - octanoic acid; C14:0 - Myristic acid; C15:0 - pentadecanoic acid; C16:0 - Palmitic acid; C18:0 - Stearic acid; C18:1n9 - Oleic acid; C18:1n7 - cis-Vaccenic acid; C20:0 - Arachidic acid; C18:3n6 - γ-Linolenic acid; C20:1n9 - 9-Eicosenoic acid; C18:3n3 - α-linolenic acid; C20:2n6 - Eicosadienoic acid; C18:3n6 - gamma linolenic acid; C20:4n6 - arachidonic acid; C20:5n3 - eico-sapentaenoic acid

Comparative analysis revealed no statistically significant differences in metabolite abundance between the transgenic and non-transgenic cowpea lines. Furthermore, the overall metabolite composition remained highly conserved across genotypes. These findings indicate that genetic transformation did not induce major metabolic perturbations within the cowpea seeds.

The results of the sub-classification of the Fatty acid methyl esters (FAME) are further presented in the **Figure 13** below. While a 60.09% saturated FAME was recorded for IT97KN, a higher percentage of 60.48 was recorded for IT97KT. Higher figure of 32.87% polyunsaturated FAME was also recorded for IT97KT in comparison to the 26.95% polyunsaturated FAME that was recorded for IT97KN. The trend was the reverse for percentage Monounsaturated FAME as the expression was higher (12.95%) for IT97KN than for IT97KT (6.64%) (**Figure 13).**

## 3.0 Discussion

Global biosafety frameworks, anchored in Article 8(g) of the United Nations Convention on Biological Diversity, underscore the need to evaluate potential unintended effects of living modified organisms, including genetically engineered (GE) crops (Secretariat of the CBD, 2000). While targeted compositional, agronomic, and phenotypic analyses have long formed the cornerstone of substantial equivalence assessments (EFSA, 2008, 2011; OECD, 2003), critics argue these approaches may miss subtle molecular perturbations arising from transgene insertion, position effects, or pleiotropy (Agapito-Tenfen et al., 2015; Corujo et al., 2018). High-throughput ‘omics technologies—proteomics and metabolomics in particular—have emerged as powerful complementary tools, enabling untargeted, systems-level characterization that can detect unintended changes and strengthen weight-of-evidence evaluations of equivalence (Christ et al., 2018; Zanon et al., 2021; Benevenuto et al., 2017).

In this study, we employed an integrated proteometabolomic approach to evaluate potential unintended molecular effects associated with the Cry1Ab transgene in the world’s first commercialized Pod Borer-Resistant (PBR) cowpea (IT97K-499-35), using its near-isogenic non-transgenic counterpart (IT97K-499-35) as the comparator. Comparative analyses of seed proteomes and metabolomes were conducted using complementary analytical platforms, including LC-MS/MS, nano-HPLC, UHPLC-MS/MS, GC-MS, and ICP-OES. The selection of the nearest isogenic line as the comparator is consistent with internationally recognized biosafety assessment frameworks, including those of AHTEG (2011) and EFSA (2011), and follows established practices in previous omics-based evaluations of genetically engineered crops (Albo et al., 2007; Zolla et al., 2008; Barros et al., 2010; Zhao et al., 2013; Benevenuto et al., 2017; Yanhua et al., 2017; Zhang et al., 2018; Zanon et al., 2021).

### 3.1 Cry1Ab Transformation Produced Limited Proteomic Perturbation

One of the most important observations of the present study is the remarkably small number of proteins that exhibited significant differential abundance following transformation. Volcano plot analyses revealed that the overwhelming majority of proteins remained unchanged between IT97KT and IT97KN, with only a limited subset exceeding the thresholds for differential expression (Figure 1; Tables 1a and 1b). Proteomic analyses of transgenic rice, maize, wheat, pea, and tobacco have similarly reported that only a small proportion of proteins differ significantly from conventional comparators and that most observed differences fall within the range of natural biological variability (Albo et al., 2007; Zolla et al., 2008; Rocco et al., 2008;). In contrast, Mesnage et al. (2016) reported more extensive proteomic alterations in NK603 maize. The discrepancy between those findings and the present study likely reflects differences in host genotype, transformation event, transgene construct, tissue type, environmental conditions, and analytical approaches.

### 3.2 Differentially Abundant Proteins Primarily Reflect Stress Adaptation and Cellular Homeostasis

Most differentially abundant proteins identified in IT97KT were associated with stress adaptation, antioxidant defense, storage functions, redox regulation, and cellular maintenance. These included members of the LEA family, thioredoxin 1, superoxide dismutase, formate dehydrogenase, mucin-like proteins, and RmlC-like jelly roll fold proteins (Table 1a, Figure 9a). Among these proteins, the LEA family deserves particular attention because it represented one of the most consistent responses observed in the transgenic line. LEA proteins are highly conserved stress-associated proteins that accumulate during dehydration, salinity stress, temperature extremes, and osmotic stress conditions (Hunault and Jaspard, 2010; Magwanga et al., 2018).

Similarly, the increased abundance of thioredoxin 1 and iron superoxide dismutase suggests maintenance of cellular redox homeostasis. Both proteins play central roles in protecting cells against reactive oxygen species and preserving protein functionality under stressful environmental conditions. Redox-regulatory proteins are among the most responsive components of plant stress-defense systems, and modest changes in their abundance are frequently observed during developmental transitions and environmental adaptation. Their presence among the differentially abundant proteins therefore supports the view that the observed changes represent physiological fine-tuning rather than adverse consequences of transformation.

### 3.3 Environmental Adaptation Pathways Remained Largely Conserved

A novel contribution of this study was the development of a stress-responsive and adaptation-related protein framework for the systematic evaluation of proteins implicated in environmental adaptation and ecological response processes. Climate change, emerging pathogens, and environmental disturbances impose growing challenges on agricultural systems, underscoring the importance of molecular mechanisms that support plant adaptation to environmental stress (Ghatak et al., 2017). The identification of 37 adaptation-associated proteins provided a targeted framework for evaluating whether Cry1Ab insertion was associated with changes in proteins involved in stress response and environmental adaptation.

Despite the large number of proteins assigned to this category, only four proteins exhibited significant abundance differences between IT97KT and IT97KN, namely LEA1, LEA4, CPRD22, and Bg7S (Table 2). The overwhelming majority of adaptation-related proteins remained unchanged, including heat shock proteins, glutaredoxins, lipid-transfer proteins, universal stress proteins, peroxiredoxins, dehydrins, and metallothioneins. This observation aligns with several proteomic studies on *Bt* crops, which generally report minimal unintended effects on stress-response pathways following transgene insertion (Gong et al., 2013; Liu et al., 2015; Wang et al., 2015). For instance, research by Wang et al. (2015) on *Bt*-transgenic and non-transgenic cotton leaves reported that the insertion of *Bt* genes caused only limited proteomic changes, with most stress-related proteins remaining stable. More recently, Nagamalla et al. (2021) found in a comparative proteomic analysis of *Bt*-cotton and non-*Bt* cotton under drought stress that the transgenic event itself induced relatively few baseline proteomic alterations in stress-defense proteins compared to non-transgenic counterparts, primarily with changes appearing under combined drought conditions. Similarly, Guan et al. (2023) reported in *Bt* Cry1Ac transgenic oilseed rape that transgenic manipulation caused only minor proteomic variations (mostly in energy-related proteins) with no biologically significant disruption to overall stress-response pathways.

This high degree of conservation is particularly noteworthy. These protein families constitute core components of the plant stress-defense machinery, mediating responses to drought, heat, oxidative stress, and other environmental challenges (Candat et al., 2014; Hibshman et al., 2021). Comparable stability in stress-related proteomes has been observed in other *Bt* transgenic systems (e.g., Cry1Ab/Cry1Ac rice and cotton), where most adaptation-related proteins showed no substantial alterations, supporting the conclusion that *Bt* events typically preserve baseline environmental resilience (Ricroch et al., 2011; Wang et al., 2015).

However, contrasts exist with prior research. While this study observed limited changes (primarily in specific LEA proteins), other investigations have documented broader proteomic reprogramming in stress-response networks. For example, Guan et al. (2023) reported differential expression of multiple proteins, including stress-related pathways, in *Bt* Cry1Ac transgenic oilseed rape. Similarly, Zanon et al. (2021) identified unintended alterations in energy metabolism, redox homeostasis, and expression of pathogenesis-related proteins in field-grown *Bt* maize. Such differences likely reflect context-specific factors such as the host genotype, transformation event, tissue type, or environmental conditions rather than a universal effect of Cry1Ab expression.

### 3.4 Ecological Risk-Associated Proteins Remain Largely Unchanged

The ecological significance of genetically engineered crops extends beyond the transformed plant to potential interactions with non-target organisms and broader ecosystem dynamics. For this reason, proteins with known or suspected roles in ecological interactions were evaluated separately. This category included pathogenesis-related proteins, defensins, osmotin-like proteins, carbonic anhydrase, and other stress-response proteins (Table 3).

Among these, carbonic anhydrase 2-like showed a statistically significant difference in abundance between IT97KT and IT97KN. In contrast, defensins, pathogenesis-related proteins, osmotin-like proteins, and other defense-associated proteins remained largely unchanged. This observation aligns with numerous studies on *Bt* crops demonstrating minimal unintended effects on ecological risk-related proteins. For instance, Wang et al. (2015) reported only limited proteomic changes in *Bt*-transgenic cotton leaves, with most defense-related proteins remaining stable. More recently, Guan et al. (2023) found that *Bt* Cry1Ac transgenic oilseed rape exhibited only minor proteomic variations, with no biologically meaningful disruption to stress-response or defense pathways. Similarly, Nagamalla et al. (2021) observed relatively few baseline alterations in defense proteins in *Bt*-cotton under stress conditions.

Meta-analyses and field studies of *Bt* crops have consistently shown negligible impacts on non-target organisms and associated molecular pathways, supporting the conclusion that modern *Bt* events generally preserve ecological safety (Wolfenbarger et al., 2008; Koch et al., 2015). However, contrasts exist with some prior research. While the present study detected only isolated changes, certain investigations have reported broader proteomic shifts in defense-related proteins (including PR proteins and osmotin-like proteins) under specific environmental conditions or in different genetic backgrounds (Guan et al., 2023; Benevenuto et al., 2017). These discrepancies are likely attributable to context-specific factors such as the host genotype, transformation method, or experimental conditions rather than a universal consequence of Cry1Ab expression.

### 3.5 Functional Enrichment Analyses Provide No Evidence of Pathway Disruption

Gene ontology enrichment analyses offered an important opportunity to determine whether the observed protein-level differences translated into coordinated alterations of biological pathways. While several enriched biological processes were identified among both upregulated and downregulated proteins, none remained statistically significant after false discovery rate correction (Figures 9a and 10a).

This observation represents one of the strongest pieces of evidence supporting molecular stability. In high-dimensional omics datasets, apparent enrichments frequently emerge as a consequence of stochastic variation. Multiple-testing correction is specifically designed to distinguish true biological signals from false discoveries. Therefore, the failure of enriched pathways to retain significance following FDR correction suggests that the observed enrichments are unlikely to represent systematic consequences of transgene insertion. Rather, they are more consistent with the natural variability expected within biological systems. The clustering and network analyses provide further support for this interpretation. Proteins associated with fungal defense, biotic interactions, pollen development, carbohydrate metabolism, and cell-wall processes formed coherent functional groups with strong biological relationships (Figures 9b, 9c, 10b, and 10c). This finding aligns with proteomic studies on other Bt and transgenic crops, where GO enrichment analyses similarly showed no significant pathway disruptions after FDR correction, reinforcing the conclusion of substantial equivalence at the functional level (Liu et al., 2020; Tan et al., 2019; Wang et al., 2015). For example, Liu et al. (2020) conducted iTRAQ-based proteomics on transgenic maize and reported that, although some GO terms were initially enriched, none indicated meaningful biological perturbation after statistical correction.

However, contrasts appear in certain studies under specific conditions. Some investigations have identified persistent enriched pathways related to stress or metabolism in transgenic lines, particularly when analyses were performed without stringent FDR correction or under combined environmental stresses (Benevenuto et al., 2017; Guan et al., 2023). Such differences likely stem from variations in experimental design, genetic background, or analytical stringency rather than inherent effects of Bt gene insertion.

### 3.6 Metabolomic Stability Corroborates Proteomic Stability

The metabolomic analyses independently corroborated the conclusions derived from the proteomic dataset. Although numerical differences were observed among several saturated, monounsaturated, and polyunsaturated fatty acid methyl esters, none of the detected metabolites differed significantly between IT97KT and IT97KN (Table 4; Figures 13–15).

The absence of significant metabolic perturbation is particularly important because metabolites represent the ultimate products of cellular activity and therefore provide a direct measure of biological function. If transformation had substantially altered major physiological pathways, corresponding changes would be expected within the metabolome. These findings are consistent with previous metabolomic investigations of genetically engineered crops, including Golden Rice, where observed differences generally remained within the range of conventional variation (Swamy et al., 2019).

Fatty acids play essential roles in membrane structure, energy storage, signal transduction, and stress responses (Dyer et al., 2008; Roudier et al., 2010). Consequently, the preservation of fatty acid composition provides strong evidence that fundamental metabolic processes remained stable following transformation.

## Conclusion

The present study provides the first integrated proteomic and metabolomic assessment of the commercially released transgenic pod borer-resistant cowpea (IT97KT) expressing the Cry1Ab gene. Comparative analyses revealed a high degree of molecular similarity between the transgenic line and its non-transgenic isoline (IT97KN), with only a limited number of differentially abundant proteins and no significant alterations in metabolite composition or abundance. Proteins associated with environmental adaptation, stress responses, and ecological interactions remained largely conserved, while multivariate analyses consistently demonstrated substantial overlap between the molecular profiles of the two genotypes. These findings indicate that Cry1Ab insertion did not result in widespread unintended perturbations of the cowpea seed proteome or metabolome. The limited molecular differences observed were comparable to the natural biological variation expected among closely related crop genotypes and were not associated with major disruptions of pathways linked to environmental adaptation or ecological risk. Collectively, the results provide strong molecular evidence supporting the substantial equivalence of transgenic pod borer-resistant cowpea and its conventional counterpart.

Given the importance of cowpea to food security and sustainable agriculture in sub-Saharan Africa, these findings contribute valuable scientific evidence to ongoing biosafety evaluations of genetically modified crops. More broadly, the study demonstrates the utility of integrated omics approaches for detecting unintended molecular changes and informing evidence-based environmental risk assessment. Future studies incorporating multi-environment evaluations and additional ecological endpoints will further strengthen our understanding of the long-term environmental performance of transgenic cowpea under diverse agroecological conditions.

